## Supplemental Figures for "Cell-autonomous timing drives the vertebrate segmentation clock’s wave pattern"

For

**Supplementary Materials**

Figures 1 – 4 supplement figures

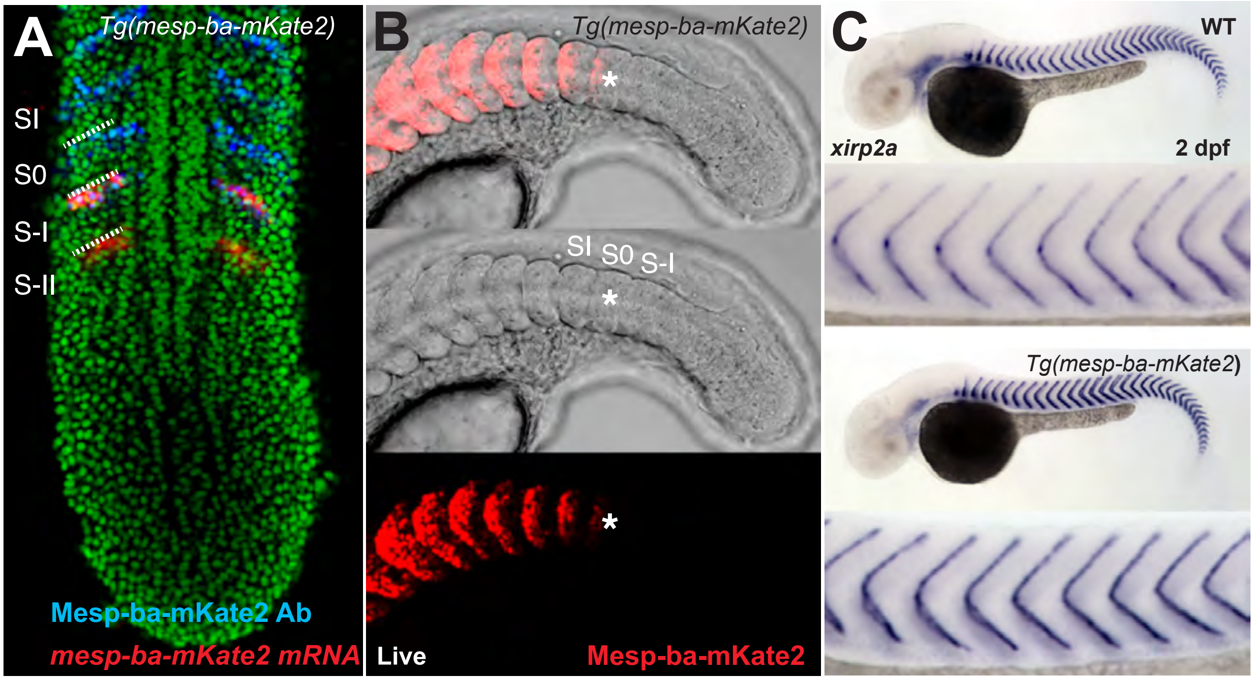


**Figure 1 – figure supplement 1. Mesp-ba-mKate2 transgenic expression and detection.** **(A)** Detection of *mesp-ba-mKate2* mRNA (red, in situ hybridization) and Mesp-ba-mKate2 (blue, anti-mKate2 antibody) in a fixed *Tg(mesp-ba-mKate2)* 10 somite-staged embryo (nuclei labelled green). *mesp-ba-mKate2* was detected only in the rostral half of pre-segments (S-II and S-I) in the anterior PSM as expected from endogenous *mesp-ba* expression (Cutty et al., 2012). Mesp-ba-mKate2 was first detected by antibodies to mKate2 in the rostral half of S-I, where it persisted in the newly forming somite (S0) and formed somites (SI). **(B)** Mesp-ba-mKate2 signal was first detected in live *Tg(mesp-ba-mKate2)* embryos within the rostral half of S0 (*), after which it remained in the rostral half of the formed somites (21 somite-stage embryo). **(C)** Boundary formation was normal as detected by in situ hybridization for the boundary marker *xirp2a* in *Tg(mesp-ba-mKate2)* embryos compared to wildtype (WT) at 2 days post-fertilization (dpf).

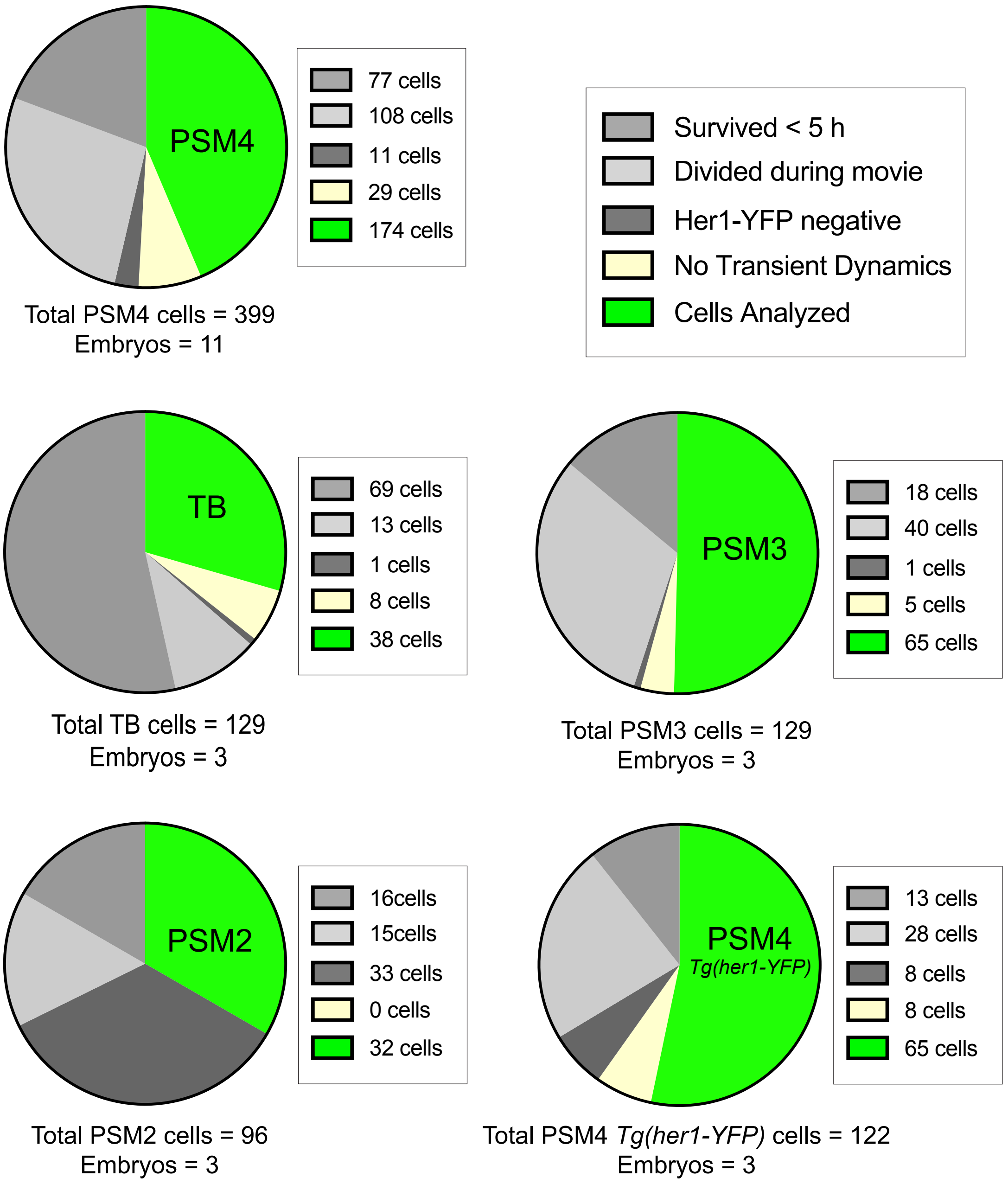


**Figure 1 – supplement figure 2. Cell culture analysis criteria.**

Dissociated cells originating from different anteroposterior positions (PSM2, PSM3, PSM4 and TB) in *Tg(her1-YFP;mesp-ba-mKate2)* embryos, or PSM4 from *Tg(her1-YFP)* control embryos. Analysis criteria was as follows: 1) Single cells alone in the field of view were selected at the start of imaging; 2) Cells dying before 5 hours post-dissociation were excluded from analysis; 3) Cells that survived > 5 h but divided during the movie were excluded; 4) From the remaining cells, those not expressing Her1-YFP or failing to oscillate and arrest before cell death were excluded (No Transient Dynamics); 5) Transient Dynamics were then analyzed in the remaining cells. Note that PSM2 cells listed as Her1-YFP negative may have already arrested before imaging started.

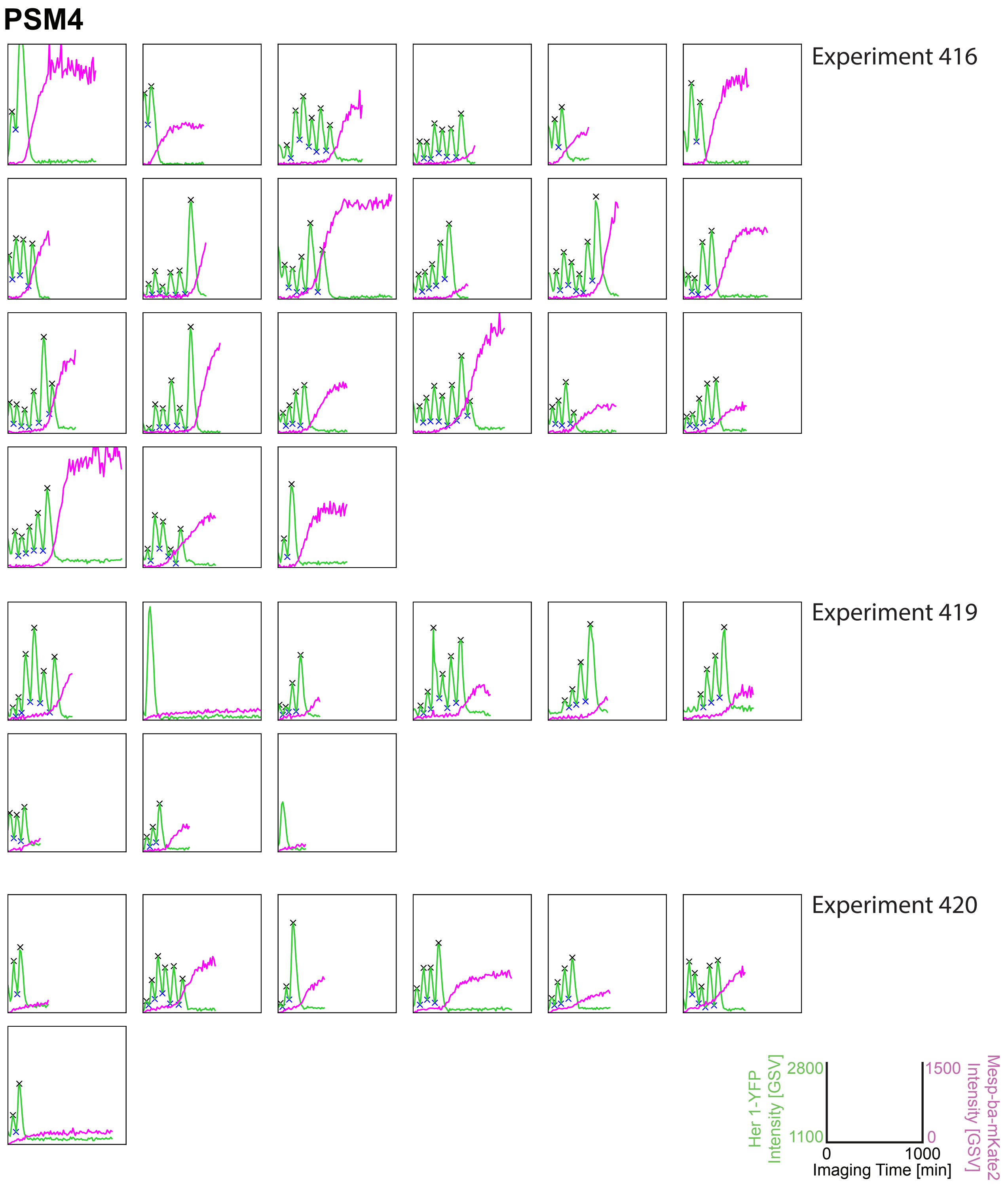


**Figure 1 – supplemental figure 3 part 1. Her1-YFP and Mesp-ba-mKate2 intensity traces from PSM4 cells in culture.** Her1-YFP and Mesp-ba-mKate2 intensity traces from single cells with the peaks and troughs marked (X) and experiment number. (N = 11 experiments, n = 174 cells).

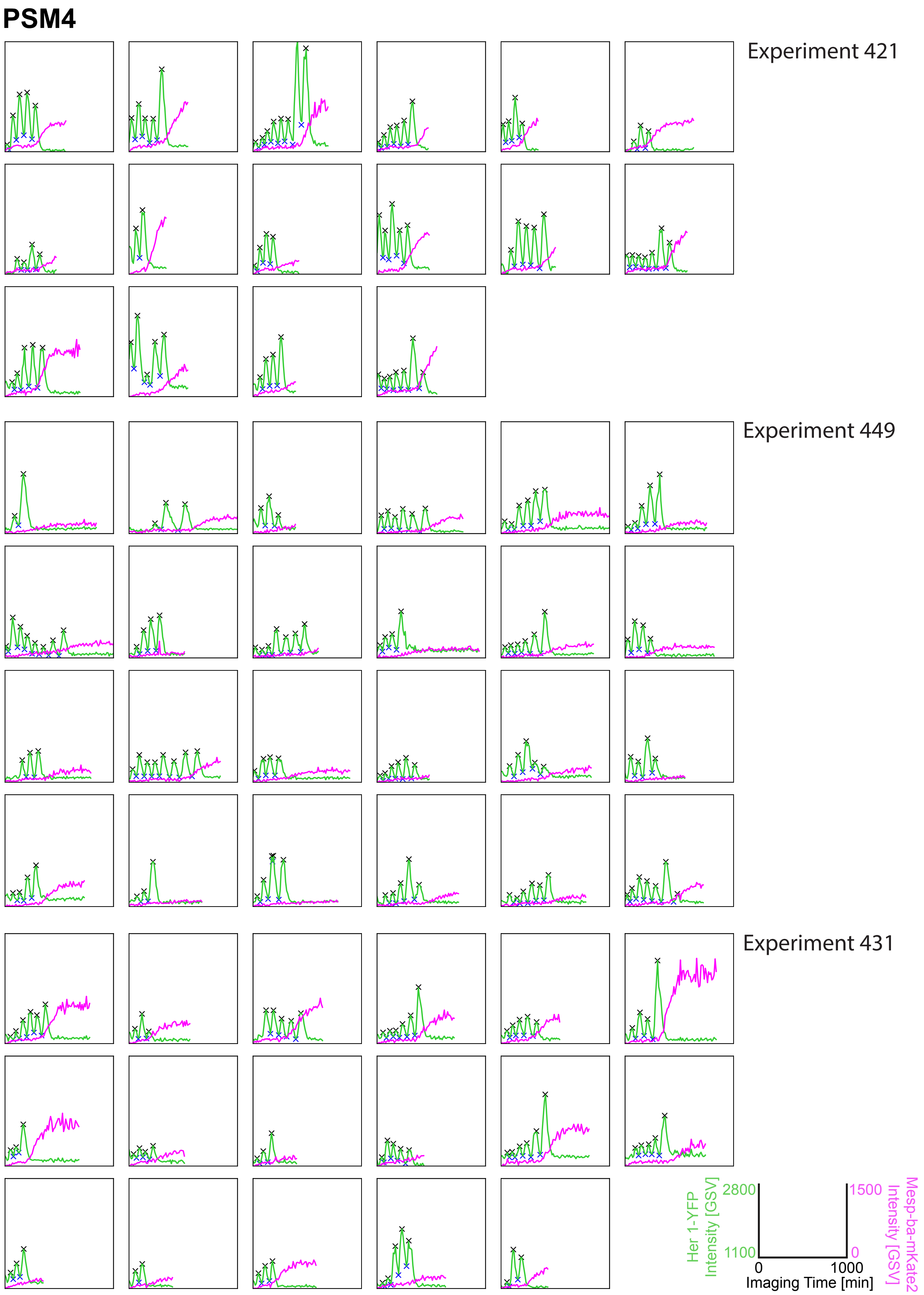


**Figure 1 – supplemental figure 3 part 2. Her1-YFP and Mesp-ba-mKate2 intensity traces from PSM4 cells in culture.** Her1-YFP and Mesp-ba-mKate2 intensity traces from single cells with the peaks and troughs marked (X) and experiment number. (N = 11 experiments, n = 174 cells).

### PSM4

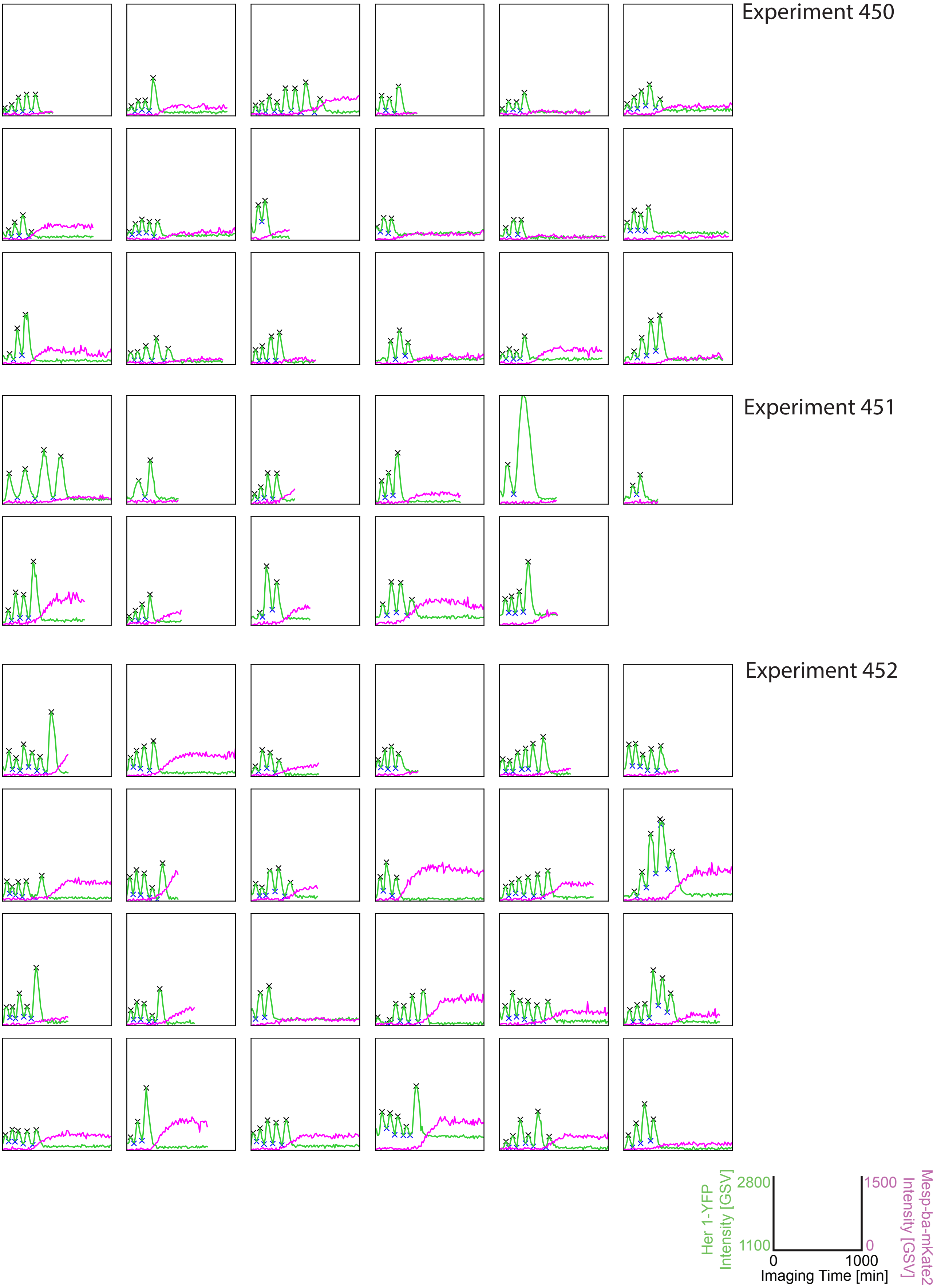


**Figure 1 – supplemental figure 3 part 3. Her1-YFP and Mesp-ba-mKate2 intensity traces from PSM4 cells in culture.** Her1-YFP and Mesp-ba-mKate2 intensity traces from single cells with the peaks and troughs marked (X) and experiment number. (N = 11 experiments, n = 174 cells).

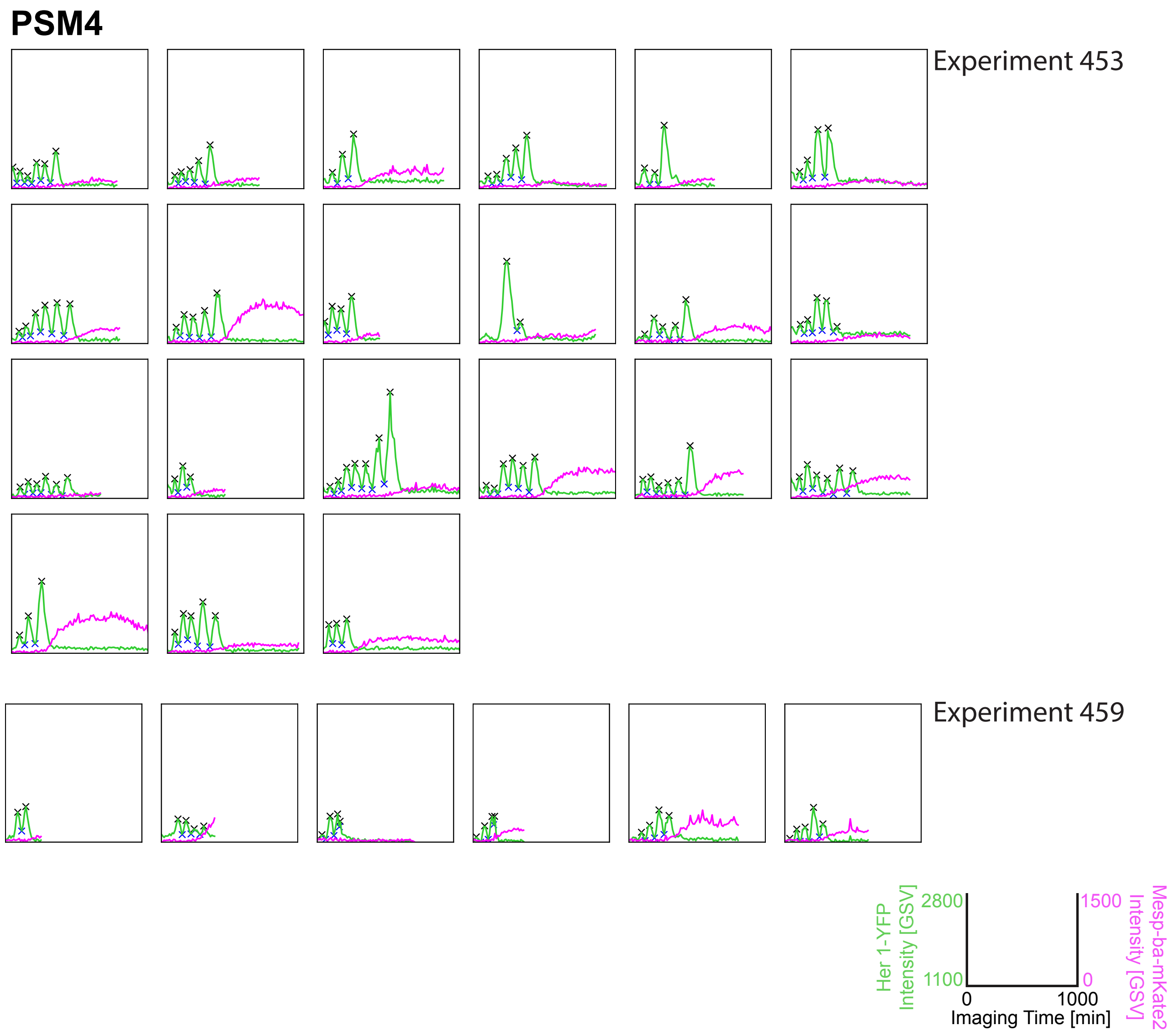


**Figure 1 – supplement figure 3 part 4. Her1-YFP and Mesp-ba-mKate2 intensity traces from PSM4 cells in culture.** Her1-YFP and Mesp-ba-mKate2 intensity traces from single cells with the peaks and troughs marked (X) and experiment number. (N = 11 experiments, n = 174 cells).

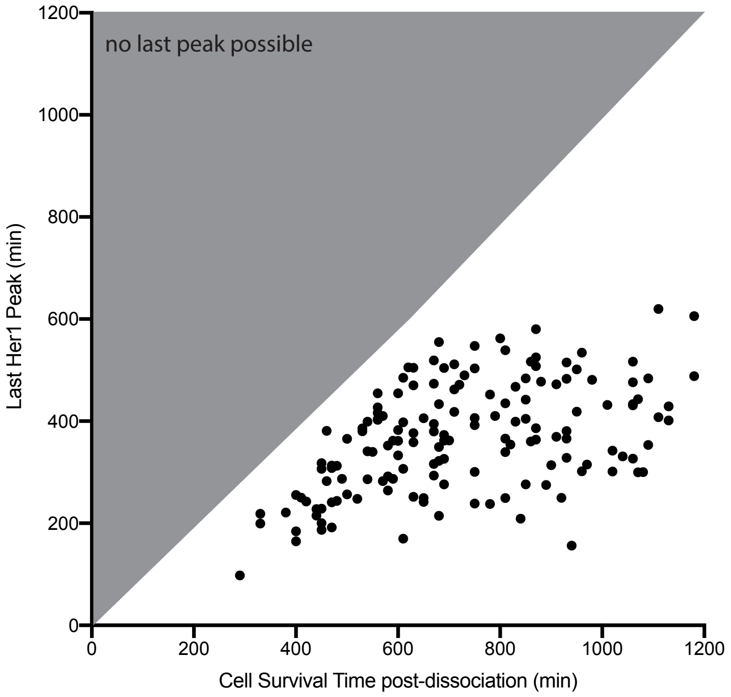


**Figure 1 – supplement figure 4. Time of cell death and oscillatory arrest in single cells.** Time of PSM4 cell death in culture and Her1-YFP last peak time in single cells. Grey triangle marks region in which a last peak is not possible because cells are already dead. Cells dying before 5 h post-dissociation were not analyzed. Cells alive at the end of imaging (>1200 min, n=20 cells) are not shown. Mean cell survival time was 787 min, SD ± 262 min.

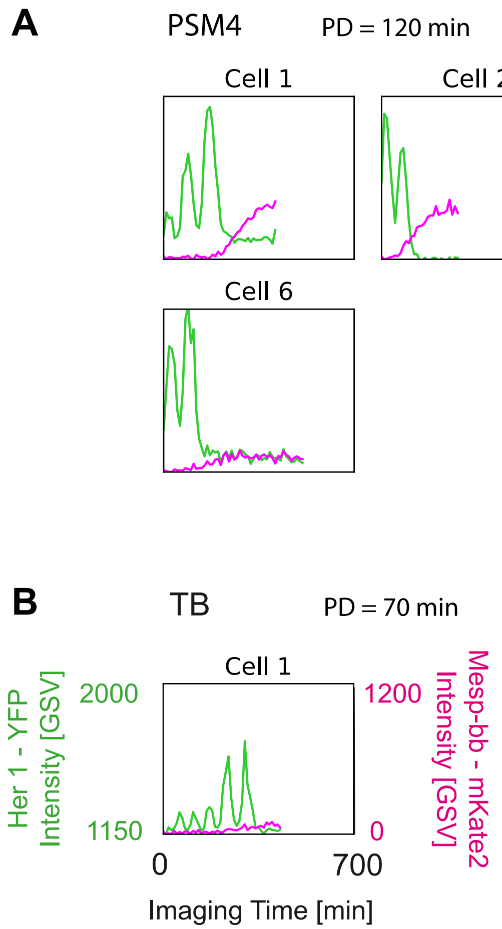


**Figure 1 – supplement figure 5. Cells isolated one-per-well reproduce autonomous behaviour.**

**(A, B)** Mesp-ba-mKate2 and Her1-YFP intensity traces in PSM4 (A) and TB (B) cells isolated one-per-well in a 24-well plate (N = 3 embryos, n = 6 PSM4 cells, n = 1 TB cell). Time of imaging start post-dissociation (PD) given. Oscillations, intensity increase and arrest in concert with Mesp-ba-mKate2 signal-onset in the isolated cells observed.

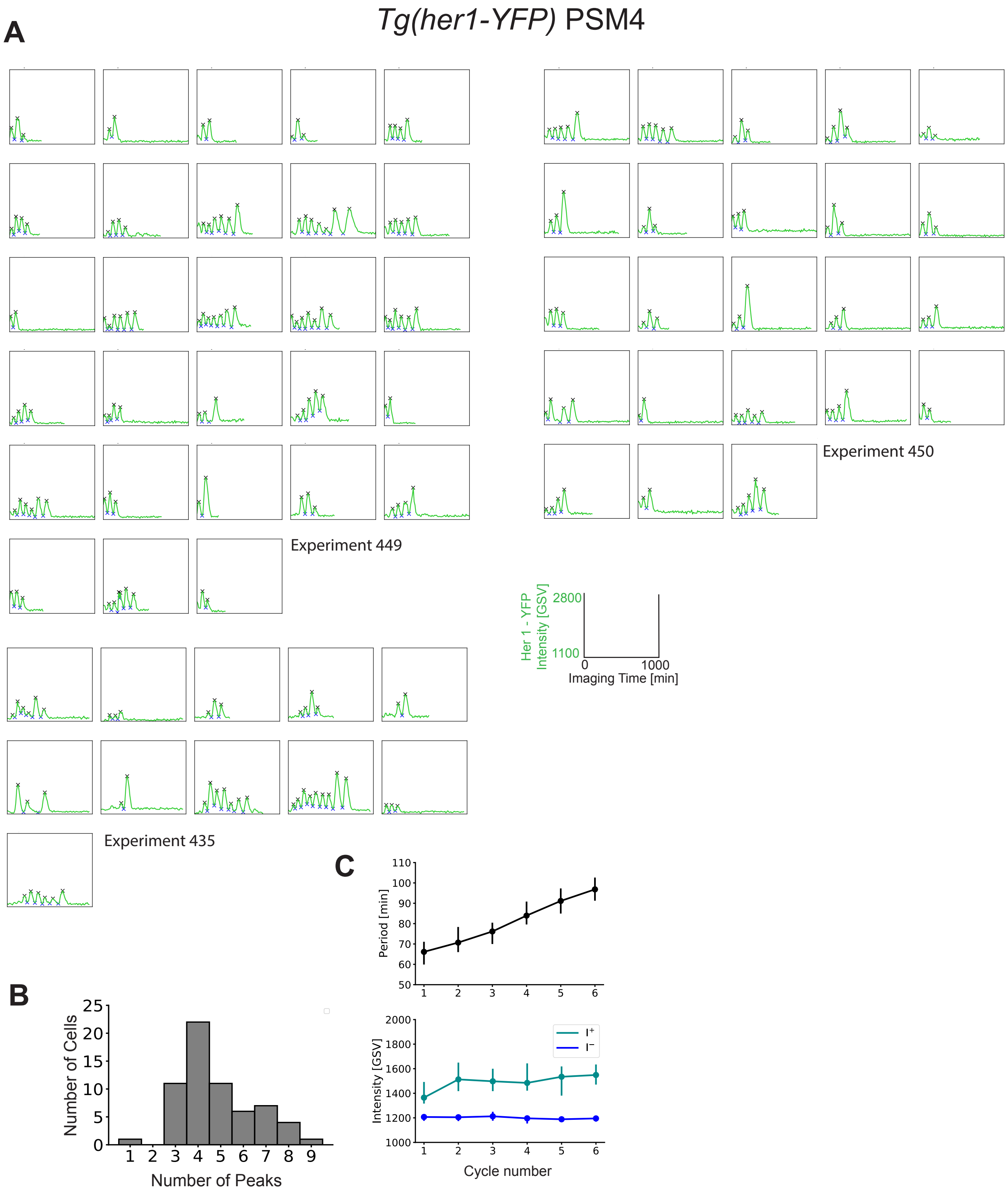


**Figure 1 - supplement figure 6. Cell-autonomous Her1-YFP dynamics do not depend on *Tg(me-sp-ba-mKate2)*.**

**(A)** Her1-YFP intensity traces in cultured PSM4 cells from embryos carrying only *Tg(her1-YFP)* (N = 3 embryos; n = 63 cells) that were cultured in parallel to PSM4 from *Tg(her1-YFP;me-sp-ba-mKate2)* embryos (experiments 449 and 450 in Figure 1 - supplement figure 3). **(B)** Number of Her1-YFP peaks. **(C)** Her1-YFP intensity traces show on average lengthing period, increasing peak intensity (I+) and constant trough intensity (I-) over successive cycles.

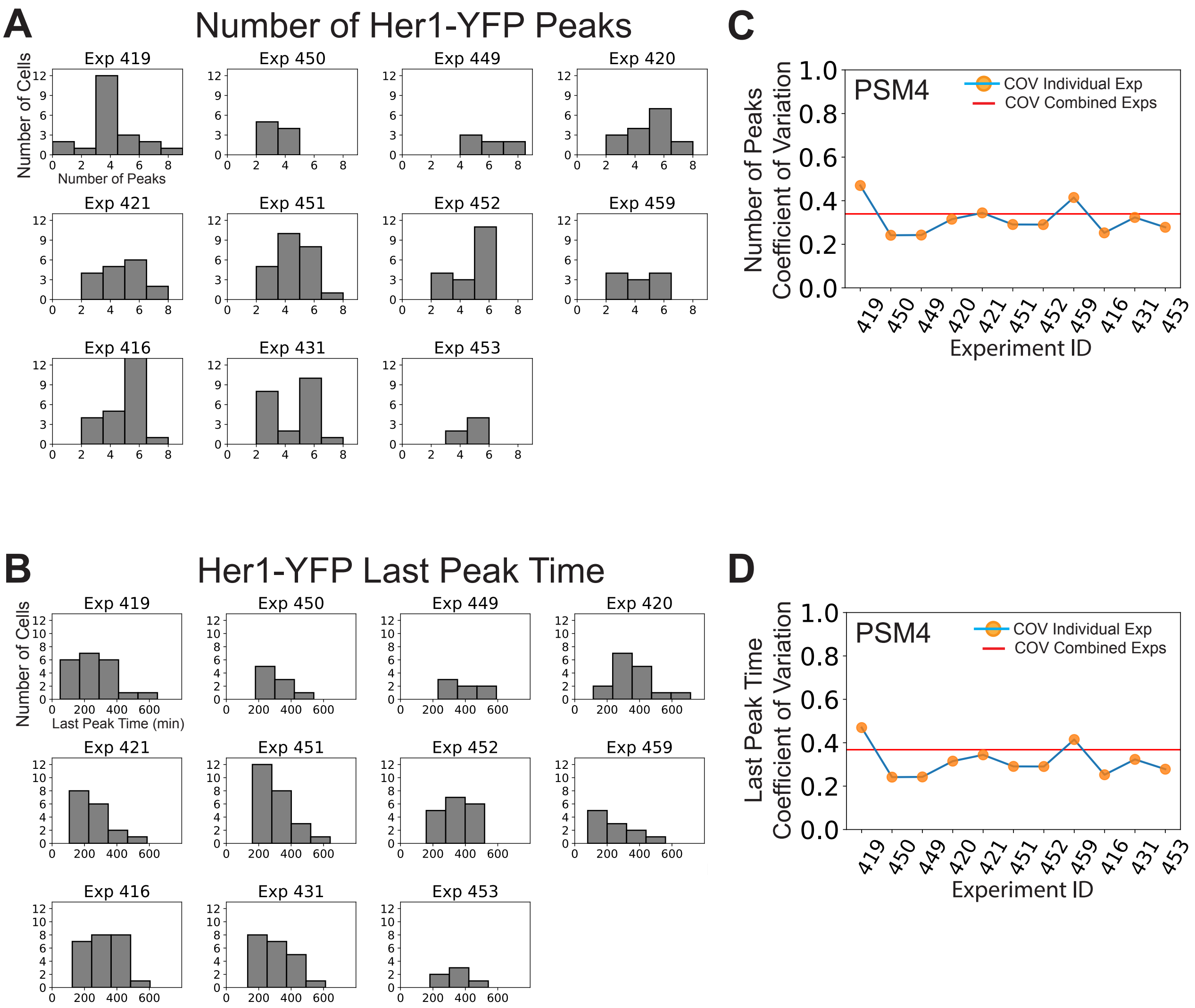


**Figure 1 – supplement figure 7. Variability in cell-autonomous timing of oscillatory arrest is not due solely to inter-experimental differences.**

PSM4 cell data in Figure 1 was pooled from 11 different experiments, each with one embryo. **(A)** The distribution of the numbers of Her1-YFP peaks generated by cells is shown for each experiment. **(B)** The distribution of the time of the Her1-YFP last peak in cells for each experiment. **(C, D)** Coefficient of Variation (COV) for each individual experiment (orange points, blue line) and mean COV for the combined set of experiments (red line) for both numbers of peaks (C) and last peak time (D).

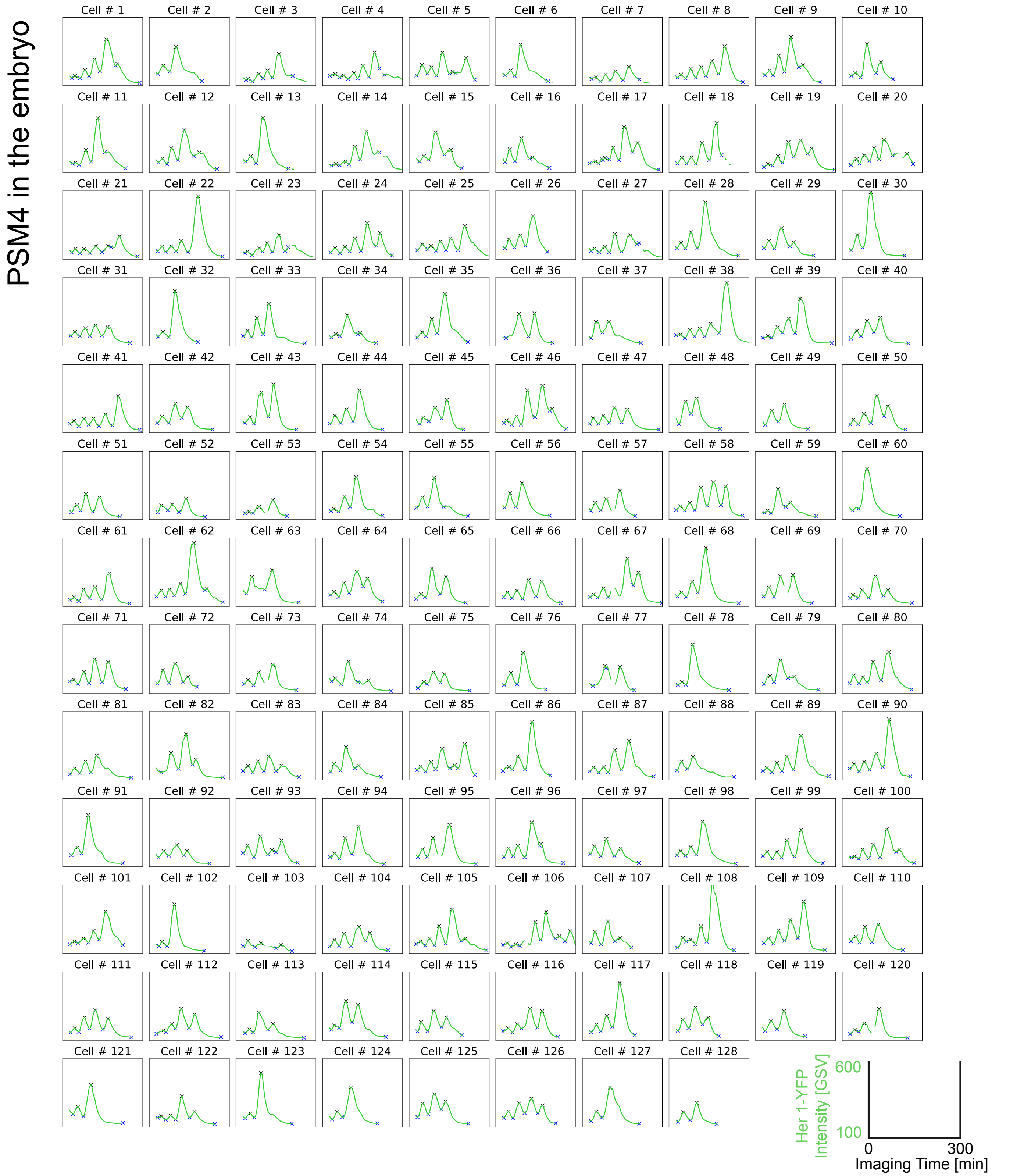


**Figure 1 - supplement figure 8. Her1-YFP intensity traces from PSM4 cells tracked in the embryo until somite formation.**

Cells were selected within PSM4 of 15 somite-staged *Tg(her1-YFP;h2b-mCherry)* embryos, then tracked until somite formation as shown in Figure 1G,H (N=2 embryos, n=128 cells). Her1-YFP intensity traces with peaks and troughs (X).

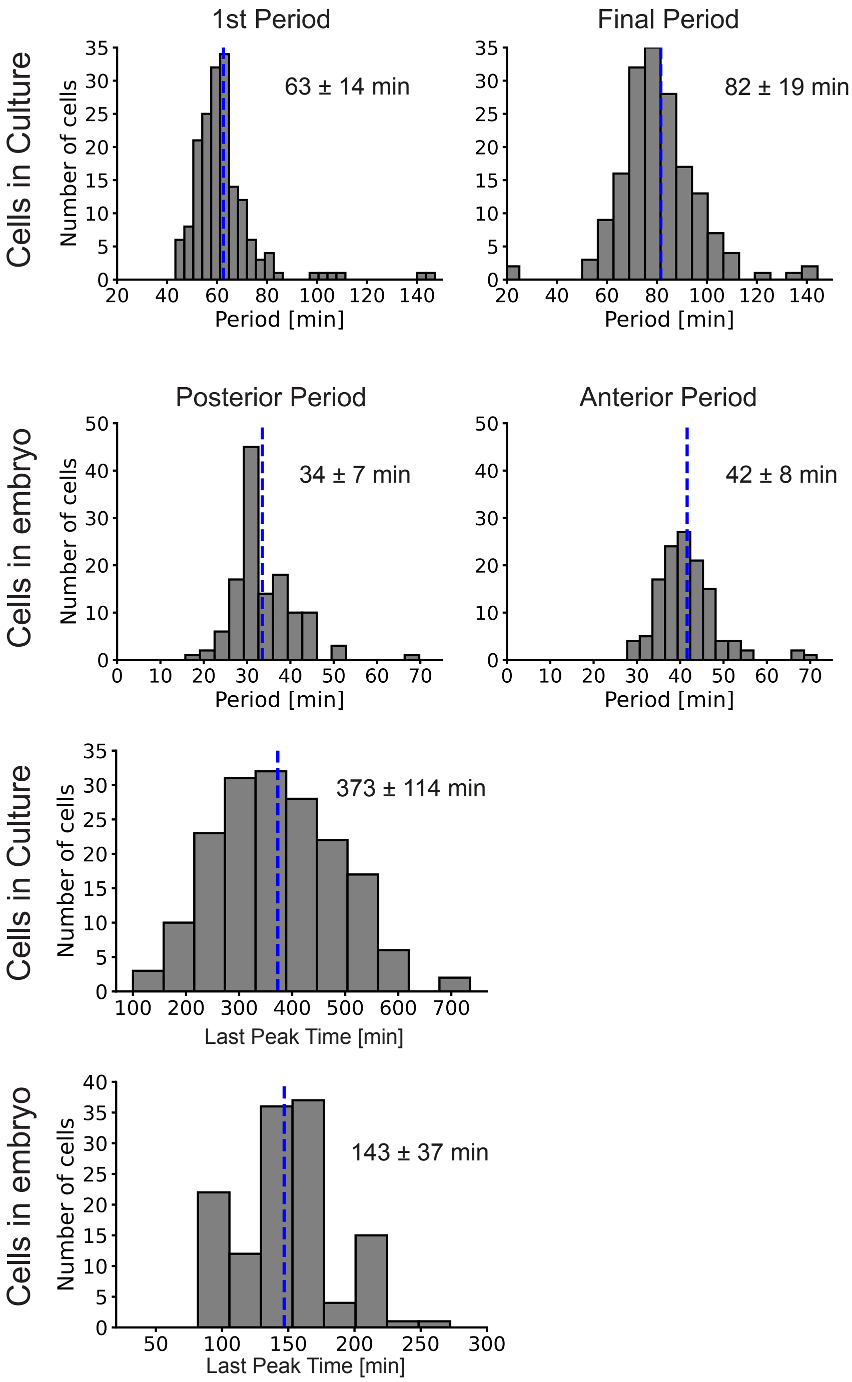


**Figure 1 – supplement figure 9. Lengthening of period and cell-autonomous transient dynamics in culture.**

**(A)** First and final periods (mean ± SD in blue) of Her1-YFP oscillations in PSM4 cells in culture. **(B)** Posterior- and Anterior-most periods of Her1-YFP in PSM4 cells tracked in the embryo. **(C-D)** Time of the Her1-YFP last peak in PSM4 cells in culture (C) and in the embryo (D). The mean of the last peak time in culture is more than double that in the embryo, indicating that the transient dynamics are skewed to comparatively longer times in culture.

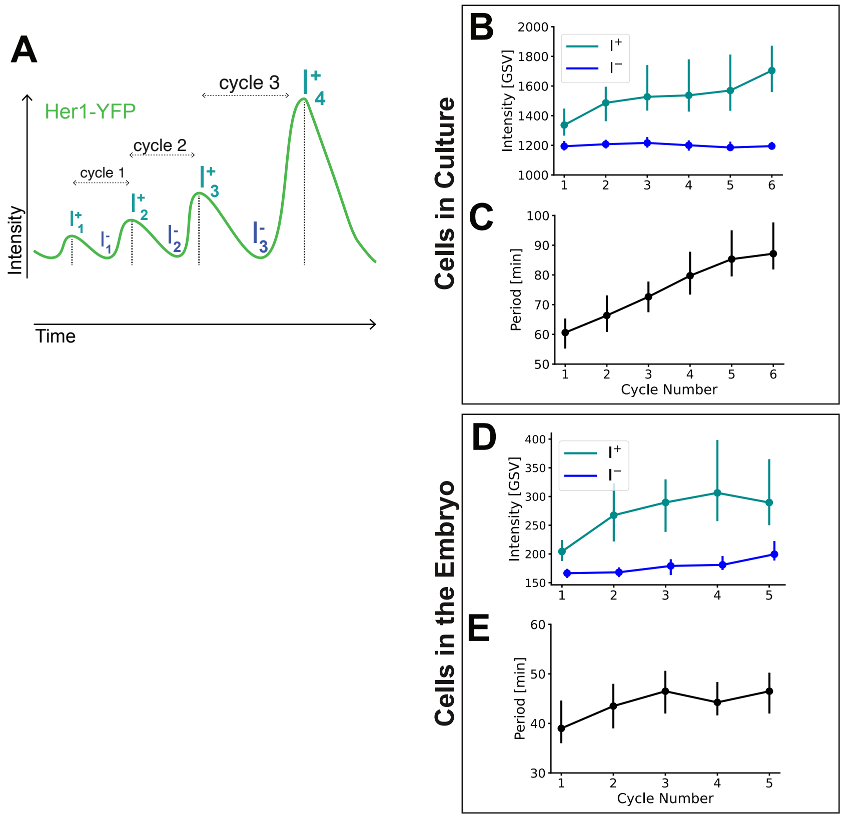


**Figure 1 - supplement figure 10.**

**(A)** Diagram of Cycle, Peak Intensities (I+) and Troughs (I-). **(B,C)** All PSM4 Her1-YFP intensity traces from cells in culture were aligned by the first peak (marked $I_1^+$ in A). Peak and trough intensity (B) and the period (C) for each cycle are given as median (circle) with 25th and 75th interquartiles (vertical bar). **(D, E)** All Her1-YFP intensity traces from cells in the embryo were aligned by the first peak, then peak and trough intensity (D) and the period (E) for each cycle shown as in B,C.

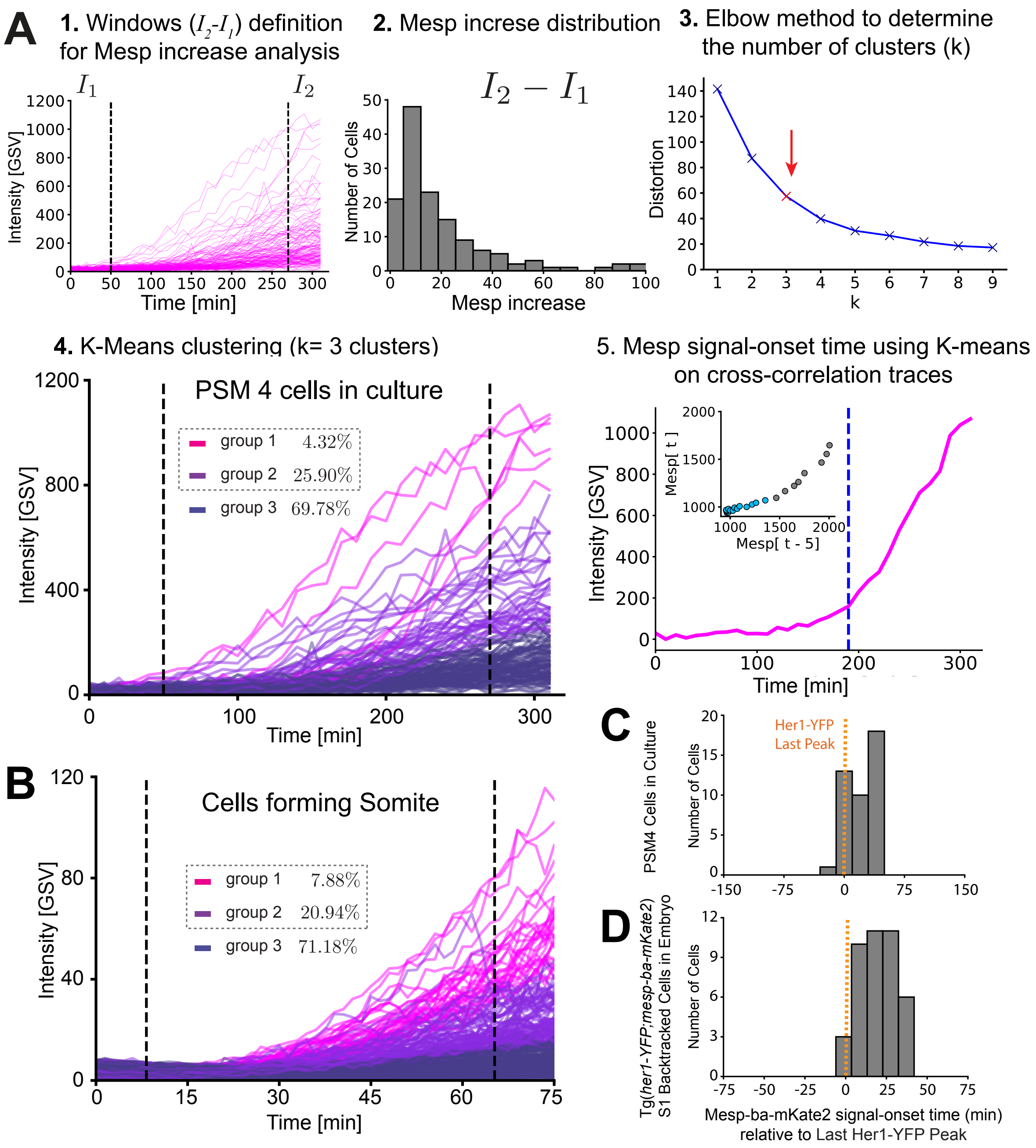


**Figure 2 - supplement figure 1. Mesp-ba-mKate2 Signal-Onset defined in individual intensity traces and timing relative to the last Her1-YFP peak.**

**(A)** Mesp-ba-mKate2 signal-onset time defined in intensity traces from PSM4 cells in culture (n = 139 PSM4 cells with sufficient trace length). Steps to define signal-onset: 1) An arbitrary window was defined across the intensity traces, then average intensity within each window was calculated. 2) Mesp-ba-mKate2 increase was obtained by subtracting the first intensity window ($I_1$) from the second ($I_2$), shown as a distribution. 3) Because intensity trace profiles varied between cells, we used the elbow method to identify the number of clusters. 4) Mesp-ba-mKate2 increase for each cell was then used to perform K-Means clustering using k = 3. Groups 1 and 2 are cells showing an obvious Mesp-ba-mKate2 signal-onset. 5) Signal-onset time was then determined in these groups using K-Means on a lag-plot of the intensity. **(B)** Mesp-ba-mKate2 signal-onset in cells forming a somite in the embryo (N = 2 somites, n = 348 pooled cells with sufficient temporal length). **(C,D)** Time of Mesp-ba-mKate2 signal-onset relative to Her1-YFP Last Peak in PSM4 cells in culture (n =42 cells) (C) and in S1 cells back-tracked in an embryo carrying both *mesp-ba-mKate2* and *her1-YFP* transgenes (n = 41 cells).

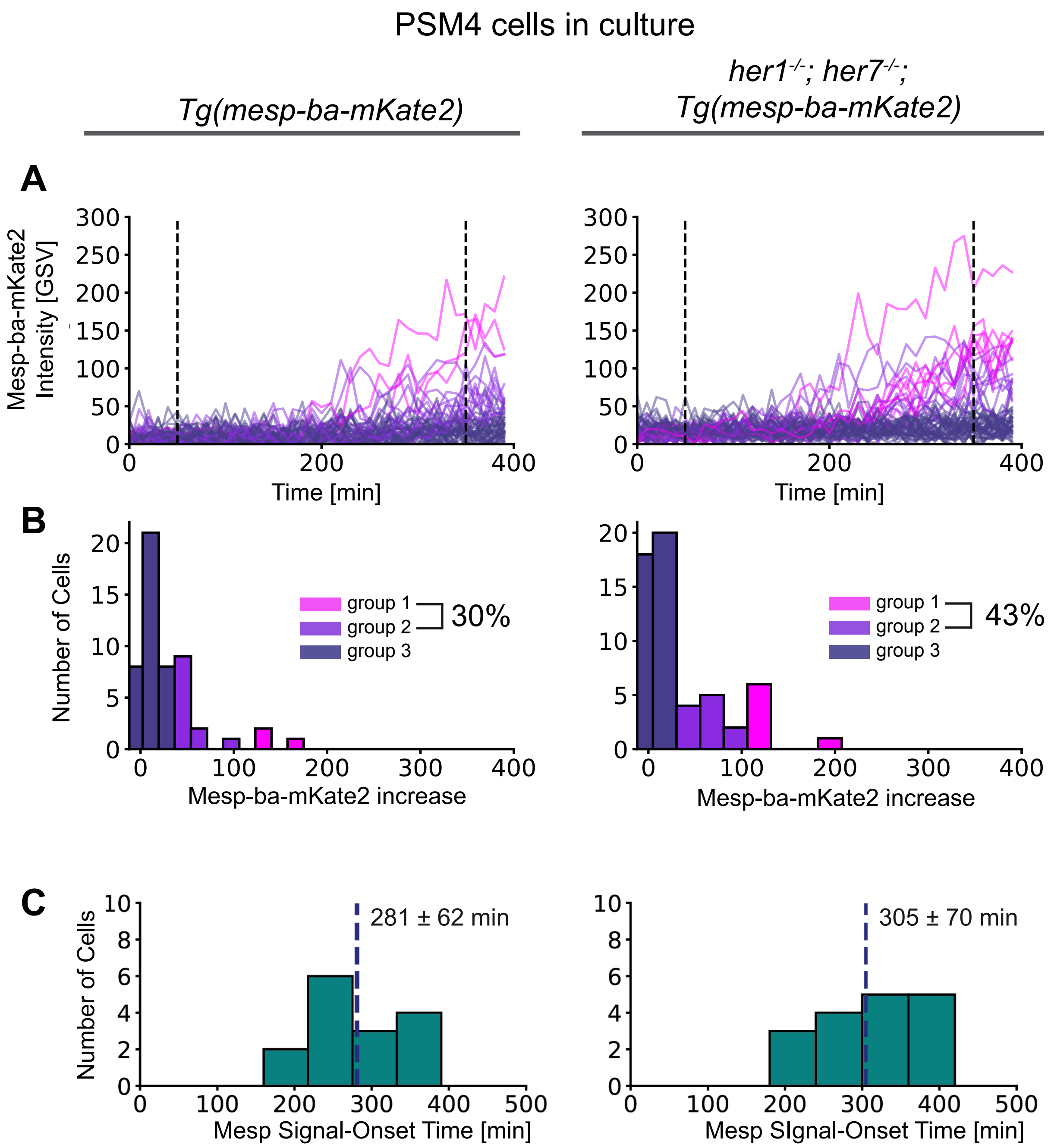


**Figure 2 - supplement figure 2. Mesp-ba-mKate2 signal-onset in her1$^{-/-}$;her7$^{-/-}$ cells with disabled segmentation clock.**

**(A to C)** PSM4 cells were cultured in parallel from her1$^{-/-}$;her7$^{-/-}$ mutant embryos carrying *Tg(mesp-ba-mKate2)* (N = 2 embryos, n = 72 cells), and control *Tg(mesp-ba-mKate2)* embryos (N=2 embryos, n = 78 cells). **(A)** Mesp-ba-mKate2 intensity traces of sufficient length were aligned by time (n = 52 control cells, n = 56 her1$^{-/-}$;her7$^{-/-}$ cells). **(B)** Mesp-ba-mKate2 intensity-increase clustered into groups (as described in Figure 2 - supplement figure 1). Percentage of cells with an obvious Mesp-ba-mKate2 signal-onset detected, groups 1 and 2. **(C)** Mesp-ba-mKate2 signal-onset times (mean ± SD) determined for group1 and 2 cells.

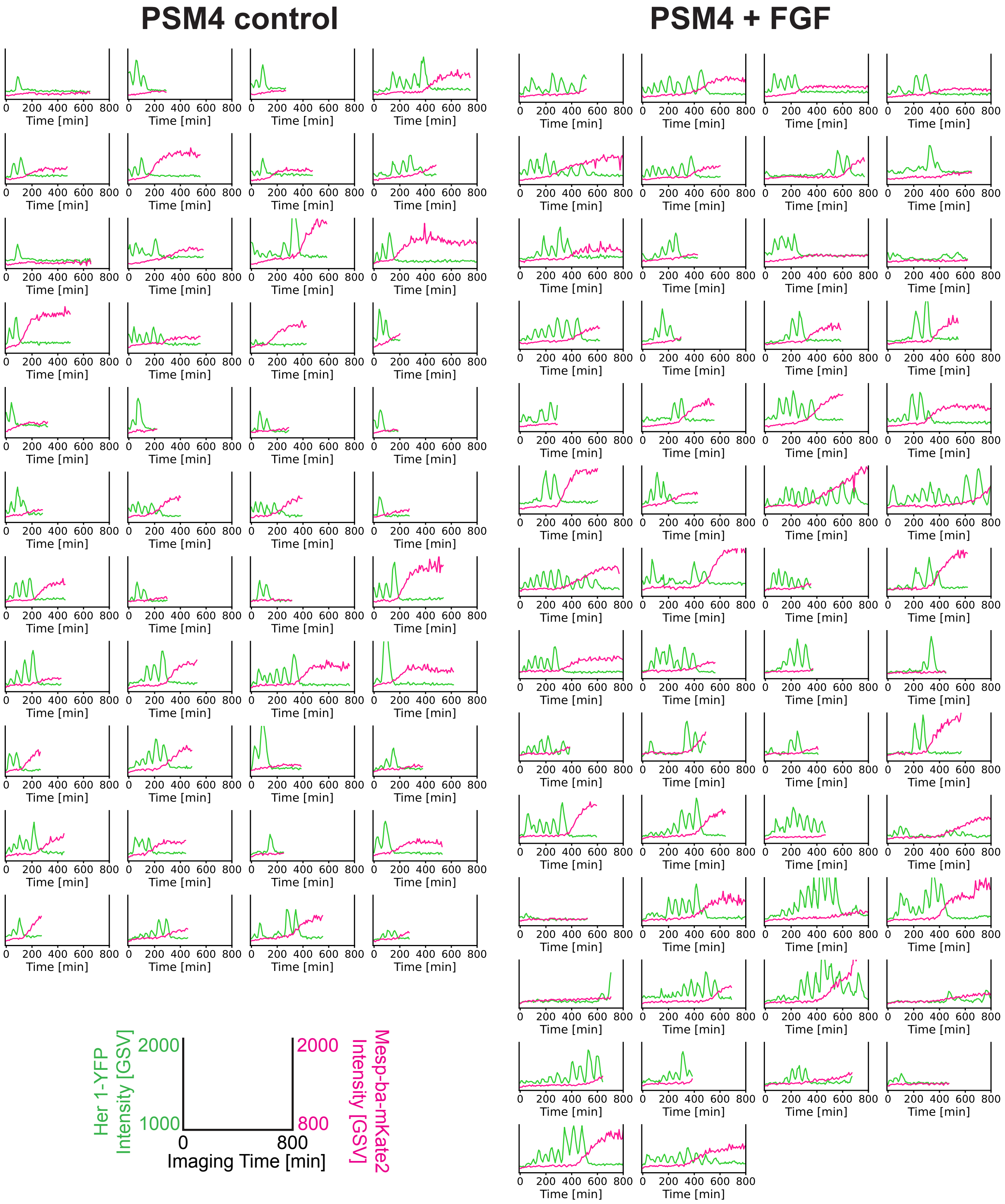


**Figure 3 - supplement 1. Her1-YFP and Mesp-ba-mKate2 cell intensity traces in PSM4 cells cultured with FGF.**

For each experiment (N = 4), dissociated cells from one or two embryos were split between two wells, then cultured +/- FGF-8b. Single oscillating cells that remained the only cell in the field of view, survived > 5h post-dissociation, did not divide, and showed transient dynamics were analyzed. Her1-YFP and Mesp-ba-mKate2 intensity traces from single cells shown (n = 44 control PSM4 cells, n = 54 PSM4 cells + FGF).

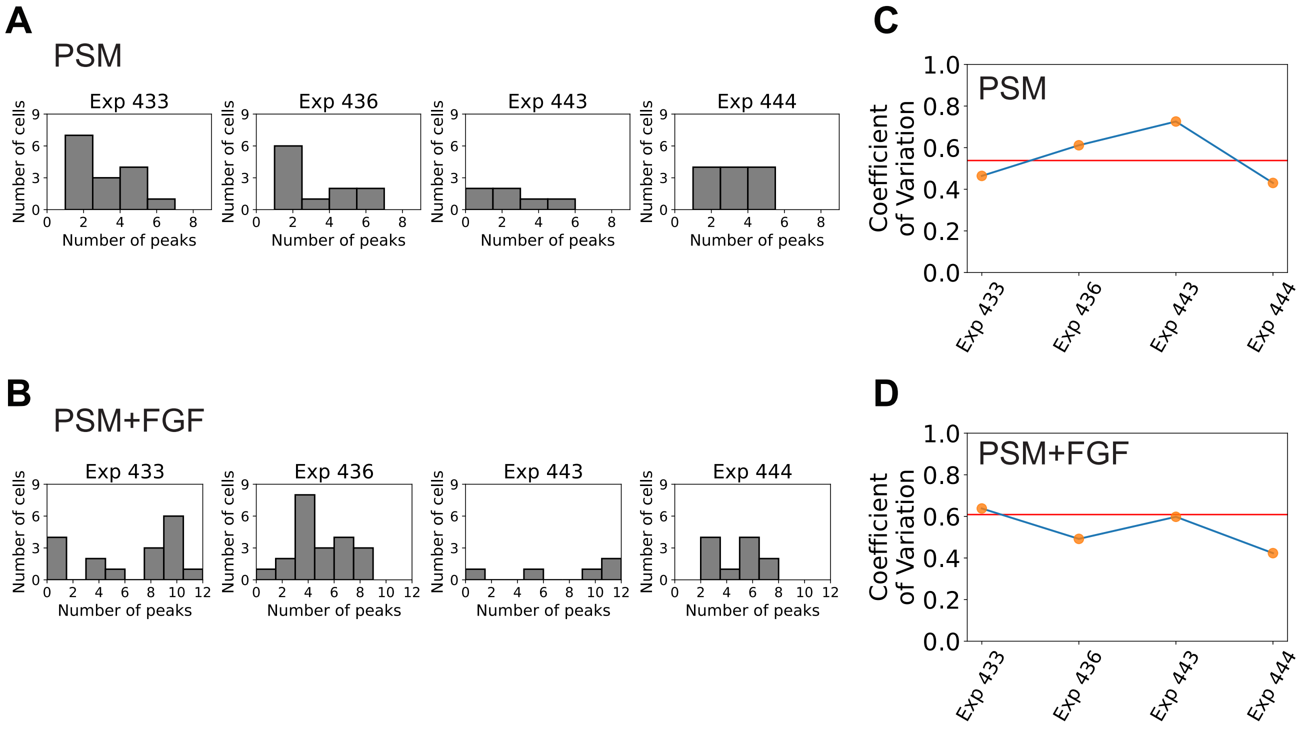


**Figure 3 - supplement figure 2 . Varibilty of cell-autonomous Her1-YFP peaks not reduced by the addition of FGF to the culture.**

**(A, B)** PSM4 and PSM4+FGF cell data was pooled from 4 different experiments. Distribution of the numbers of Her1-YFP peaks generated by individual cells shown for each experiment. **(C, D)** Coefficient of Variation (COV) for each individual experiment (orange points, blue line) and mean COV for the combined set of experiments (red line).

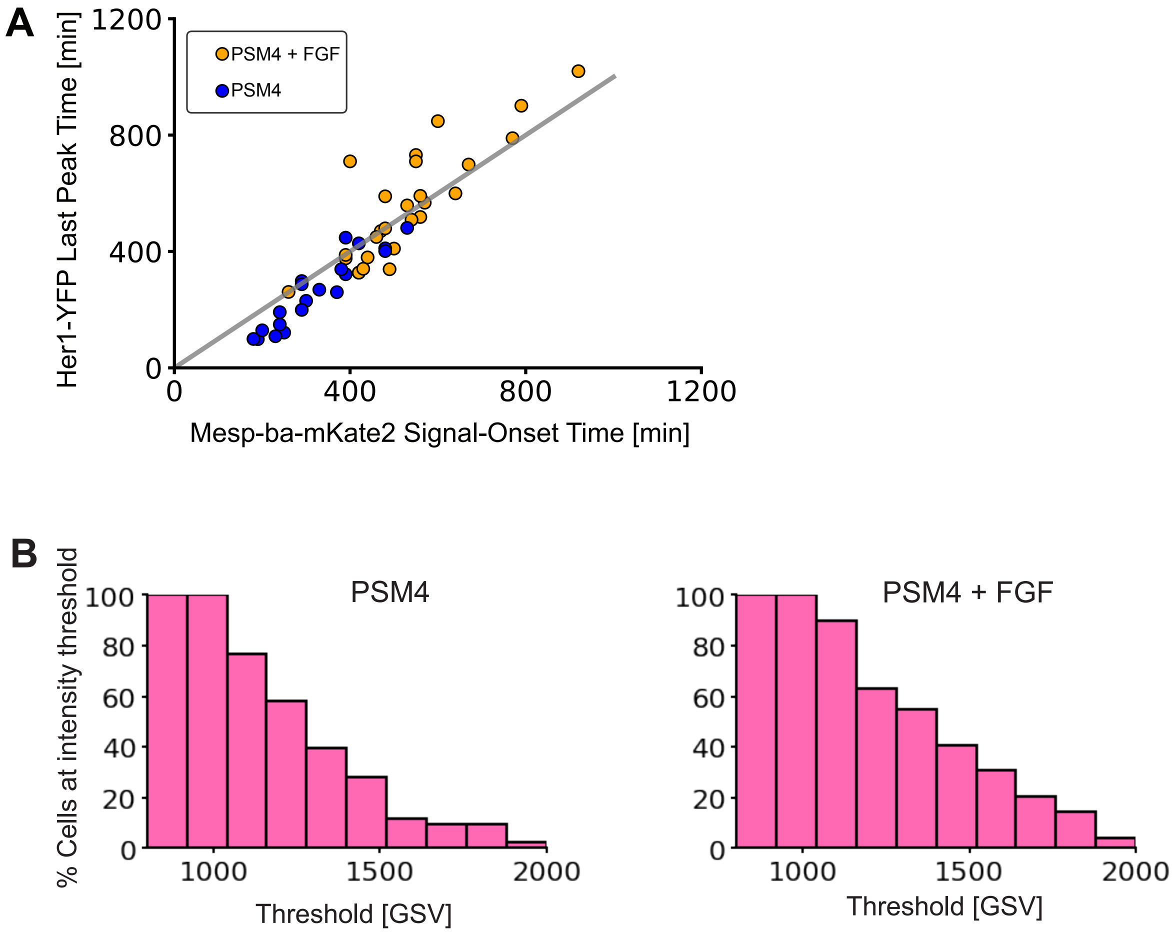


**Figure 3 - suppplement 3. Mesp-ba-mKate2 signal onset and intensity distributions in response to FGF.**

**(A)** Her1-YFP Last peak time correlated with Mesp-ba-mKate2 signal-onset time in PSM4 (blue) and PSM4 + FGF (orange). (**B)** Mesp-ba-mKate intensity shown by percent of cells reaching a range of intensity thresholds.

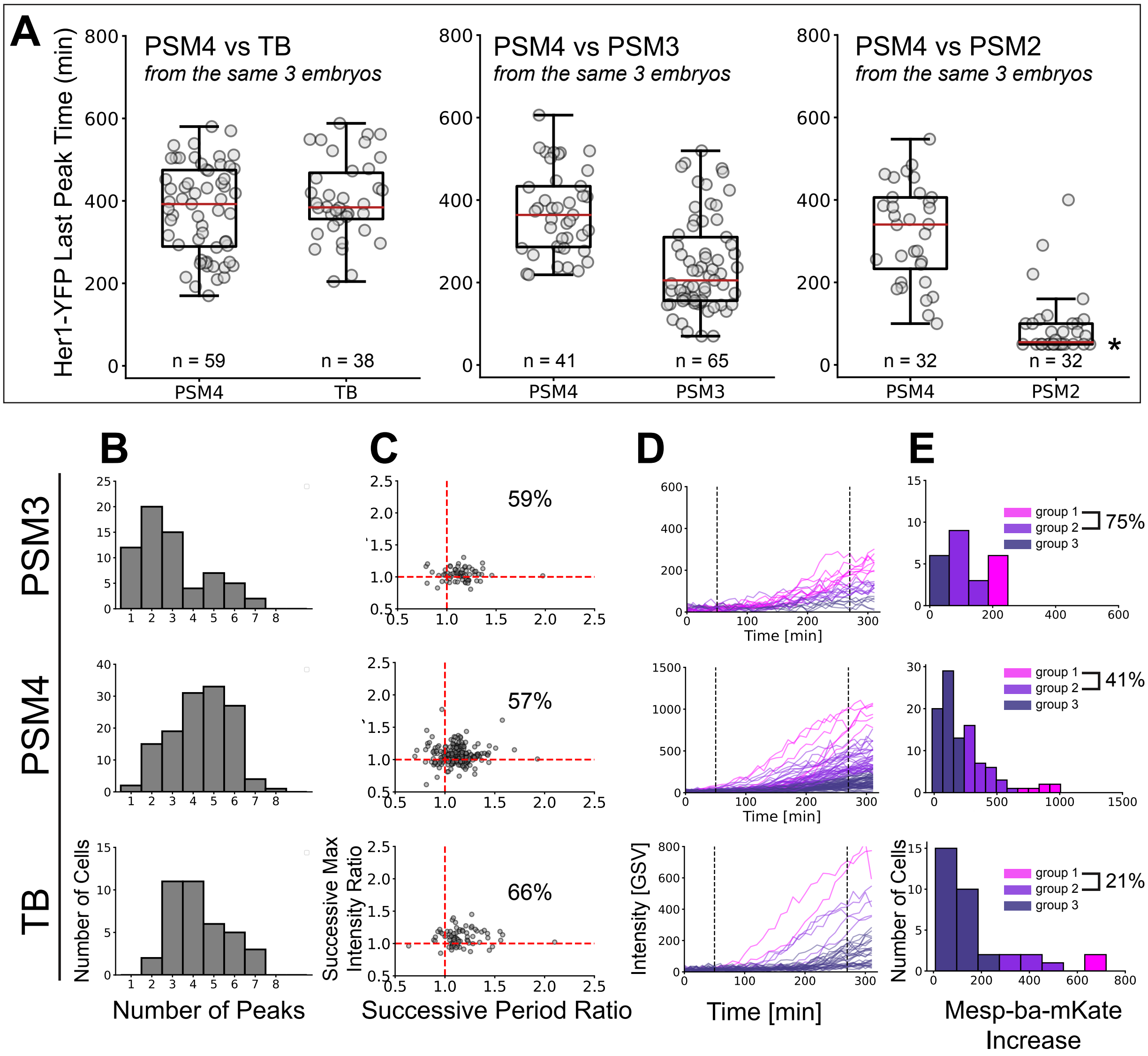


**Figure 4 – supplement 1. Cell-autonomous Her1-YFP and Mesp-ba-mKate2 dynamics in cells dissected from different anteroposterior positions.**

**(A)** Her1-YFP last peak times in cultured cells dissected from different anteroposterior positions (PSM2, PSM3, PSM4, and TB) in *Tg(her1-YFP;mesp-ba-mKate2)* embryos. PSM4 was dissected and cultured in parallel to other PSM quarters from the same embryo as an internal reference (N = 3 for each comparison). Pooled PSM4 data is shown in Figure 4. Median last peak time (red line) with interquartile box. Many PSM2 cells were in the fall of the last peak when imaging began, thus the last peak time for such cells was set to the time imaging started post-dissociation (*). **(B)** Number of Her1-YFP peaks (mean ± SD, PSM3 2.95 ± 1.65; PSM4 4.36 ± 1.44; TB 4.26 ± 1.35 peaks) **(C)** Successive period and intensity ratios. Percentage of successive cycles slowing and increasing intensity. **(D, E)** Mesp-ba-mKate2 intensity traces and distributions clustered into groups.

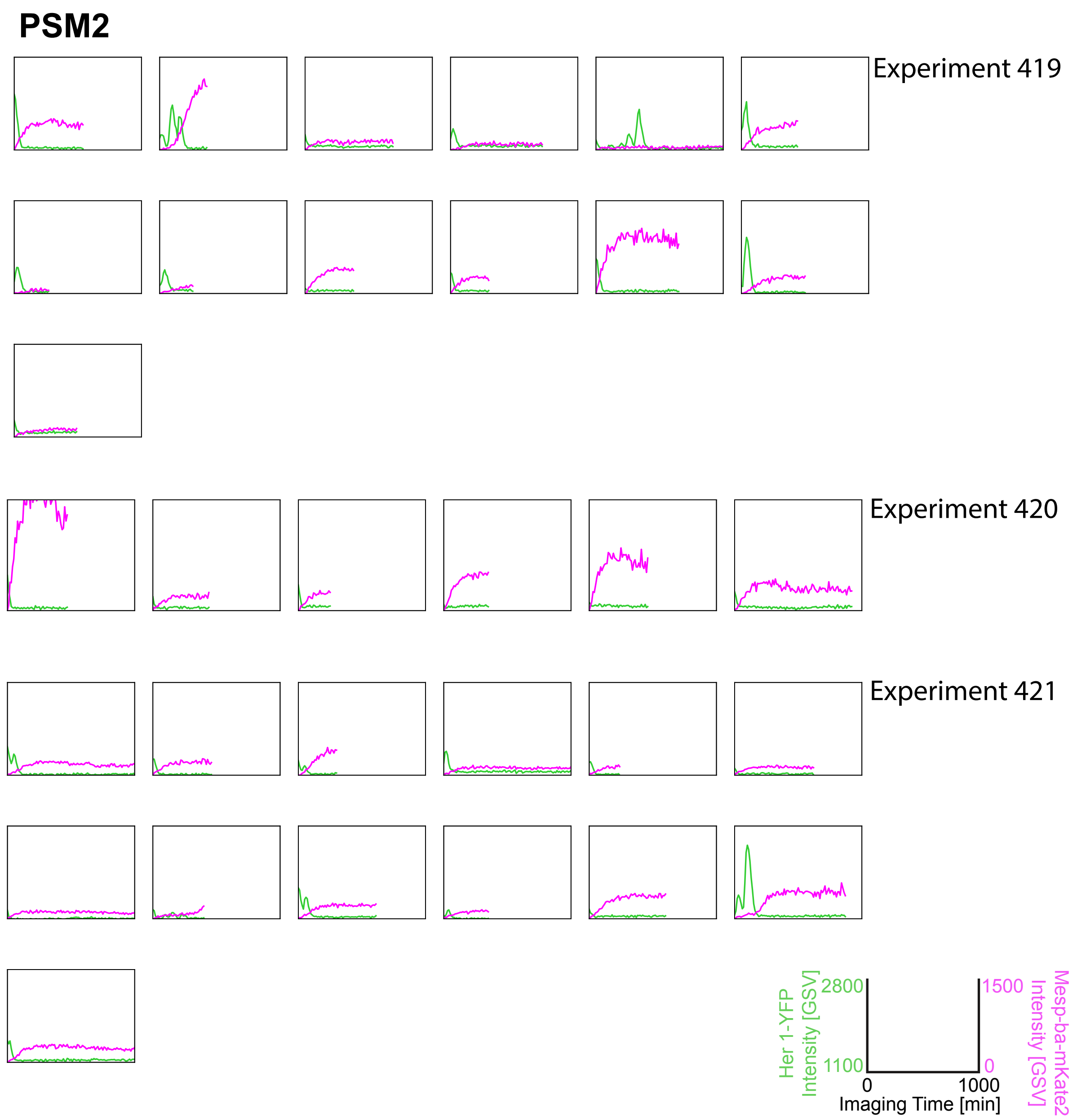


**Figure 4 - supplement figure 2. PSM2 Her1-YFP and Mesp-ba-mKate2 intensity traces.** Intensity traces from the second quarter of PSM (PSM2) (N = 3 experiments, n = 32 cells) and experiment number.

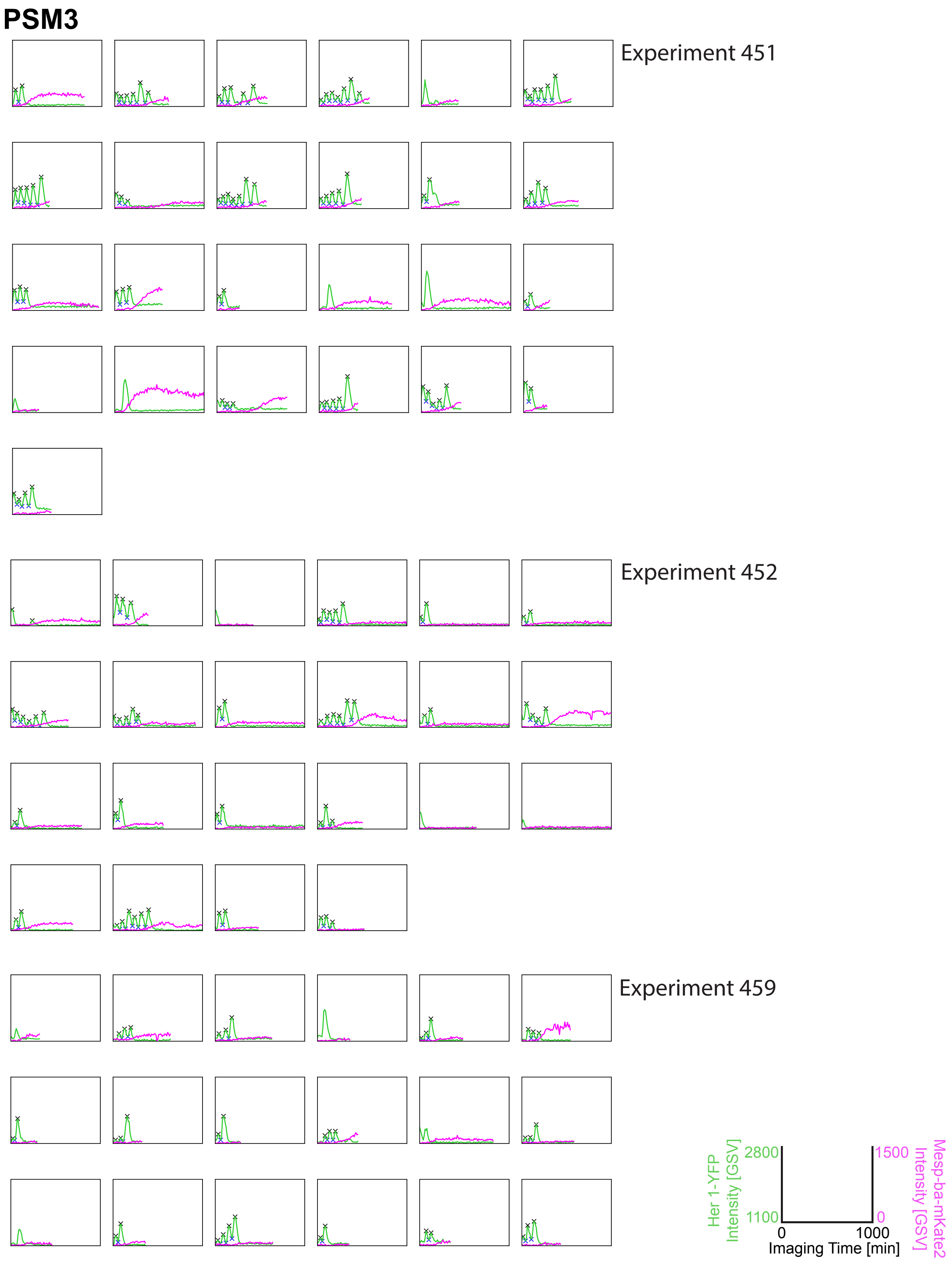


**Figure 4 – supplement figure 3. Her1-YFP and Mesp-ba-mKate2 intensity traces from PSM3 cells in culture.** Her1-YFP and Mesp-ba-mKate2 intensity traces from single cells with the peaks and troughs marked (X) and experiment number. N = 3 experiments, n = 65 cells

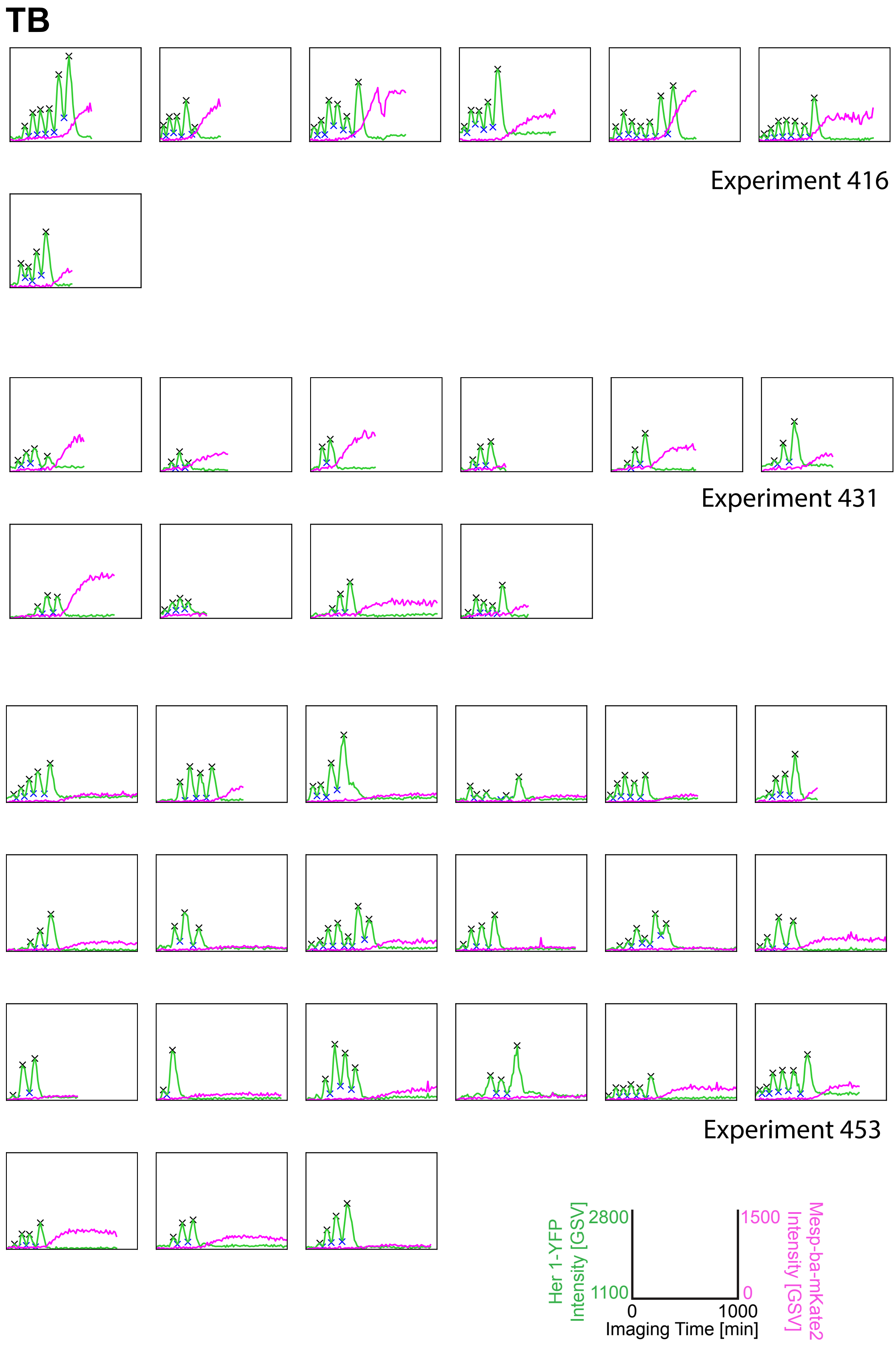


**Figure 4 - supplement figure 4. Tailbud Her1-YFP and Mesp-ba-mKate2 intensity traces.**

Intensity traces from Tailbud (TB) cells in culture (N = 3 experiments, n = 38 cells). Peak and troughs marked (X) and experiment number given.
